# Calibrated prediction of scarce adverse drug reaction labels with conditional neural processes

**DOI:** 10.1101/2024.06.07.598036

**Authors:** Miguel Garcia-Ortegon, Srijit Seal, Shantanu Singh, Andreas Bender, Sergio Bacallado

## Abstract

Adverse drug reactions (ADRs) are a major source of concern in the development of novel pharmaceuticals. ADRs may be identified in the late stages of development or even after commercialization, which may lead to failure or discontinuation after spending enormous resources on candidate molecules. Thus, predicting ADRs early in the process could help reduce costs by avoiding future failures. However, due to the low number of drugs approved, the amount of historical datapoints on ADRs is limited, which makes their prediction challenging for traditional chemoinformatics methods. Interestingly, each approved drug may have been annotated for hundreds of ADRs, which opens the door to framing ADR prediction as a multi-task or meta-learning problem. In this work, we adopt a meta-learning approach to ADR prediction by applying conditional neural processes (CNPs) to the publicly available Side Effect Resource (SIDER). Our results suggest that CNPs are competitive against single-task baselines even when trained on sparse datasets with missing labels. Furthermore, we find that their predictions are well-calibrated. Finally, we evaluate their performance on ADRs associated to different physiological systems and confirm good predictions across organ classes. Our findings suggest that meta-learning strategies may be beneficial for data-limited clinical endpoints like ADRs.

## 1 Introduction

A major challenge to discovering novel therapeutics is the emergence of unexpected adverse drug reactions (ADRs), also known as side effects, at later stages of the development pipeline. Lead-stage ADRs can lead to the discontinuation of otherwise promising compounds after significant resources have been invested. In addition, unexpected ADRs can lead to the removal of drugs during postmarketing surveillance years after approval, with huge costs to companies and immense damage to patients. Unfortunately, such occurrences are far from rare: between 1953 and 2013, over 462 compounds were reported to be withdrawn from the market, with the most common ADR being liver toxicity (Onakpoya et al., 2016). Being able to predict ADRs early in the development process could significantly reduce the cost of drug discovery by helping discard candidates that are destined to fail (Liu and Zhang, 2019).

Machine-learning (ML) models may be a helpful tool for ligand-based screening (Thomas et al., 2022), including early ADR identification (Lee and Chen, 2019; Dey et al., 2018; Zhang et al., 2021, 2017). If a model predicts certain ADRs are associated with a candidate compound, medicinal chemists may be able to carry out tailored tests to confirm or disprove them before committing valuable resources to the candidate. However, the scarcity of side-effect labels hinders the potential applicability of data-driven methods. Only a few thousand drugs have ever been approved, so the abundance of high-confidence ADR annotations remains low. Furthermore, the fact that ADRs may be discovered years after a drug is marketed suggest that adverse-effect annotations in current datasets are likely incomplete. Stilll, even if some side effects are missing in current datasets, each approved drug may already be associated to tens or hundreds of ADRs. This suggests that ADR prediction may be framed as a multi-task or meta-learning problem. Such strategy may increase data efficiency and robustness against incomplete or sparse datasets.

Neural processes (NPs) (Garnelo et al., 2018b) are a family of neural probabilistic models for meta-learning. They can transfer information efficiently across functions in order to make predictions based on just a few labeled datapoints. Their training procedure includes random sampling of possibly non-overlapping observations from different functions, which makes them naturally suited to deal with sparse datasets. Recently, their application to molecular functions has been explored (Lee et al., 2022; Garcia-Ortegon et al., 2022; Chan et al., 2023). In this paper, we study the utilization of conditional NPs (CNPs) for ADR prediction. We evaluate CNPs on a dataset of historical side effects, the Side Effect Resource (SIDER)(Kuhn et al., 2016), using either all data available or removing a fraction of labels in order to artificially increase sparseness.

## 2 Methods

### 2.1 Conditional neural processes (CNPs)

Consider a meta-training dataset with observations of binary-valued functions *f*_1_, …, *f*_*n*_, *f*_*i*_ : 𝒳 *→* {0, 1}. In this paper, *f*_*i*_ corresponds to a particular ADR, 𝒳 represents the space of chemically feasible molecules and *x* ∈ 𝒳 refers to a single molecule represented as a fingerprint vector; in this way, binary labels 0 or 1 indicate whether a certain molecule *x* is free from or associated with a side effect *f*_*i*_. Each molecular function *f*_*i*_ is observed at a set of *O*_*i*_ input points 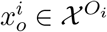, with known labels 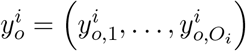, where 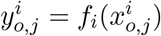.

Additionally, consider a meta-test function *f*, observed at a small set of *C context* points (*x*_*c*_, *y*_*c*_) = ((*x*_*c*,1_, *y*_*c*,1_), …, (*x*_*c,C*_, *y*_*c,C*_)). Our goal is to predict the values *y*_*t*_ of *f* at a set of *T target* locations *x*_*t*_ ∈ 𝒳^*T*^ as accurately and efficiently as possible, using the example context points (*x*_*c*_, *y*_*c*_) and the observations from the example functions *f*_*i*_, …, *f*_*n*_.

A neural process (NP) (Garnelo et al., 2018a,b) is a parametric model for meta-learning that aims to describe the predictive distribution *p* (*y*_*t*_ | *x*_*c*_, *y*_*c*_ ; *x*_*t*_) (semicolon notation is used to differentiate contexts and targets). Most NPs are designed for regression, assuming a Gaussian predictive distribution. In this paper, we use NPs for binary classification and assume a Bernouilli predictive distribution:

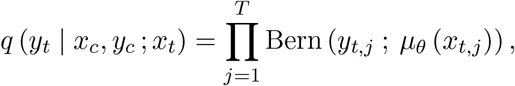

where the targets *x*_*t,j*_, …, *x*_*t,T*_ are taken to be conditionally independent. At test time, the predicted probability values for the targets are binarized, such that targets with a predicted mean 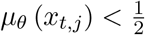 are labeled as 0, and targets with 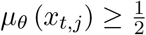 are labeled as 1. In this paper, we use the conditional NP (CNP) (Garnelo et al., 2018a), which parameterizes the predictive distribution for a point *x* in three steps. First, contexts (*x*_*c,j*_, *y*_*c,j*_) are mapped by an encoder network *h*_*θ*_ to a local datapoint representation *r*_*j*_. Second, all context encodings *r*_*j*_ are combined into a global function encoding *r* through a commutative operation, usually the sum or the mean. Commutativity guarantees invariance to contexts’ permutations. Third, a deterministic decoder network *g*_*θ*_ maps the function encoding *r* and the input location *x* to the predictive’s parameters, i.e. *µ*_*θ*_ (*x*) in a Bernoulli.

The parameters of the CNP *θ* are trained by backpropagation from the predictive log-likelihood ℒ_*θ*_(*y*_*t*_ | *x*_*c*_, *y*_*c*_ ; *x*_*t*_) = log *q*_*θ*_(*y*_*t*_ | *x*_*c*_, *y*_*c*_ ; *x*_*t*_). During meta-training, each meta-train function *f*_*i*_ is seen once every epoch, but not all observations 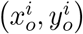 are used at each iteration. Rather, the *O*_*i*_ observations are randomly subsampled to create two disjoint sets: a context set 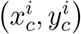 and a target set 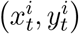, with sizes *C*_*i*_ and *T*_*i*_ respectively, *C*_*i*_ + *T*_*i*_ *≤ O*_*i*_. The predictive log-likelihood on the current targets is optimized given the current contexts. Therefore, the final objective is

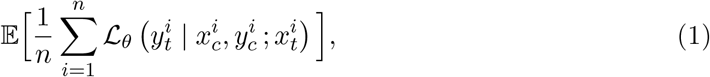

where the expectation is with respect to the random sampling procedure. *C*_*i*_ and *T*_*i*_ can themselves be stochastic: in our experiments, we sample them uniformly from [20, 150) at each iteration.

### 2.2 Baseline models

We assessed the performance of CNPs against two single-task classifiers: random forest (RF) and XGBoost (XGB). RF was implemented in scikit-learn (Pedregosa et al., 2011) with default parameters and 500 trees and XGB was implemented using the official API (XGB) with default parameters. We also compared with a random classifier which sampled labels from the label distribution of the training set.

### 2.3 Dataset and splitting

The Side Effect Resource (SIDER) (Kuhn et al., 2016) is a dataset with information about 1556 molecules (of which 1430 are drugs), 5880 ADRs and 140064 molecule-ADR pairs. In addition, ADRs are grouped into 27 system organ classes that categorize side effects by associated physiological system. In the CNP formulation, we treat ADRs as functions and molecules as datapoints. We downloaded the dataset from the SIDER website (SIDER) and retrieved isomeric SMILES strings (Weininger, 1988) from PubChem (Kim et al., 2023). Reported side effects were labeled as 1, and the absence of evidence for a side effect was labeled as 0. For our exploratory experiments, we retained side effects that had at least 100 positive labels, yielding 370 side effects and an overall rate of positive labels of *∼* 19%. In all experiments, molecules were represented as Morgan fingerprints (Morgan, 1965) of length 1024 and radius 3. We computed these using RDKit (RDKit).

The dataset was segmented randomly across functions (ADRs) and across datapoints (compounds), with a 80*/*20% split for both. We refer to these splits as *ftrain, ftest* and *dtrain, dtest* respectively (Figure 1). The CNP was meta-trained on *ftrain, dtrain* and *ftrain, dtest*. Later, during meta-testing, we applied the CNP to each ADR in *ftest*, supplying observed molecules from *ftest, dtrain* as contexts and evaluating on unobserved molecules from *ftest, dtest* as targets. For each random seed, a single CNP was trained for all functions. Baseline models were trained on *ftest, dtrain* and evaluated on *ftest, dtest*. For each random seed, separate single-task baselines were trained for each function in *ftest*.

**Figure 1:**
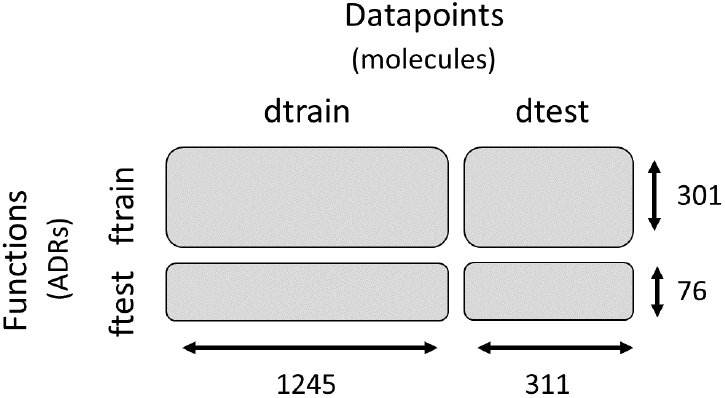
Split of SIDER across functions (ADRs) and datapoints (molecules).

To simulate learning from a sparse dataset, we performed an experiment were we randomly removed 50% of the labels of *ftrain, dtrain*, of *ftrain, dtest* and of *ftest, dtrain* of every ADR. The results of this experiment are shown in Section 3.1.

## 3 Experiments

### 3.1 Binary classification of ADRs in complete and sparse SIDER

In our first experiment, we evaluated the capability of CNPs to predict ADRs in a complete and sparse dataset. Given an ADR function and test molecules, models were required to predict a positive (1, presence of ADR) or negative (0, absence of ADR) label for each test molecule. We compared the CNP to a random classifier that predicted classes with probabilities according to their representation in the training set, to a random forest (RF) and to XGBoost (XGB) 2.2. Each model was trained 4 times with different random initializations. Table 1 shows the mean and standard error across all repetitions and across all *ftest* ADRs.

**Table 1:** Classification metrics of baseline models and CNP on *ftest, dtest*

|  | Trained on complete dataset |  |  |  |
| --- | --- | --- | --- | --- |
|  | Precision | Recall | ROC-AUC | Phi |
| Random | 0.18 (0.02) | 0.18 (0.02) | 0.50 (0.00) | -0.01 (0.01) |
| RF | 0.59 (0.02) | 0.21 (0.02) | 0.69 (0.01) | 0.25 (0.01) |
| XGB | 0.49 (0.02) | 0.25 (0.02) | 0.66 (0.01) | 0.23 (0.01) |
| CNP | **0.67 (0.03)** | **0.31 (0.02)** | **0.83 (0.01)** | **0.37 (0.02)** |

|  | Trained on 50%-sparse dataset |  |  |  |
| --- | --- | --- | --- | --- |
|  | Precision | Recall | ROC-AUC | Phi |
| Random | 0.18 (0.02) | 0.18 (0.02) | 0.50 (0.00) | -0.00 (0.01) |
| RF | 0.59 (0.03) | 0.15 (0.02) | 0.65 (0.01) | 0.19 (0.01) |
| XGB | 0.44 (0.02) | 0.21 (0.02) | 0.63 (0.01) | 0.18 (0.01) |
| CNP | **0.64 (0.02)** | **0.35 (0.02)** | **0.83 (0.01)** | **0.39 (0.02)** |

Our results suggest that the CNP beat single-task baselines in both the complete and the sparse datasets. Interestingly, the performance of single-task models degraded in the sparse version of SIDER, while CNPs maintained their original performance. This may be due to the ability of CNPs to transfer information across different functions, even if their datapoints do not fully overlap as is the case here. Performing well in datasets with missing labels may be relevant in practice because many negative labels in real ADR datasets actually missing data, in the sense that they signify a lack of evidence rather than concrete evidence of no adverse reaction.

### 3.2 Calibration of predicted probabilities

A binary classification model is said to be well calibrated if, when the model predicts a probability *p* for a certain datapoint, the true label of the datapoint turns out to be 1 a fraction *p* of times. Following Niculescu-Mizil and Caruana (2005), in order to estimate calibration we collected each model’s predictions for every molecule in *ftest, dtest* (i.e. we pooled all 76 test functions). Then, we binned molecules according to their predicted probabilities. Within each bin, we computed the average predicted probability and the fraction of molecules in the bin with true label 1, and plotted one against the other (Figure 2). The CNP and RF exhibited reasonable calibration, closely adhering to the ideal diagonal line. This is interesting because neither of these models was trained explicitly to optimize calibration; in particular, the CNP optimized a maximum predictive likelihood objective (Equation 1). In contrast, XGB showed poor calibration, with a tendency to underestimate the probability of true positives.

**Figure 2:**
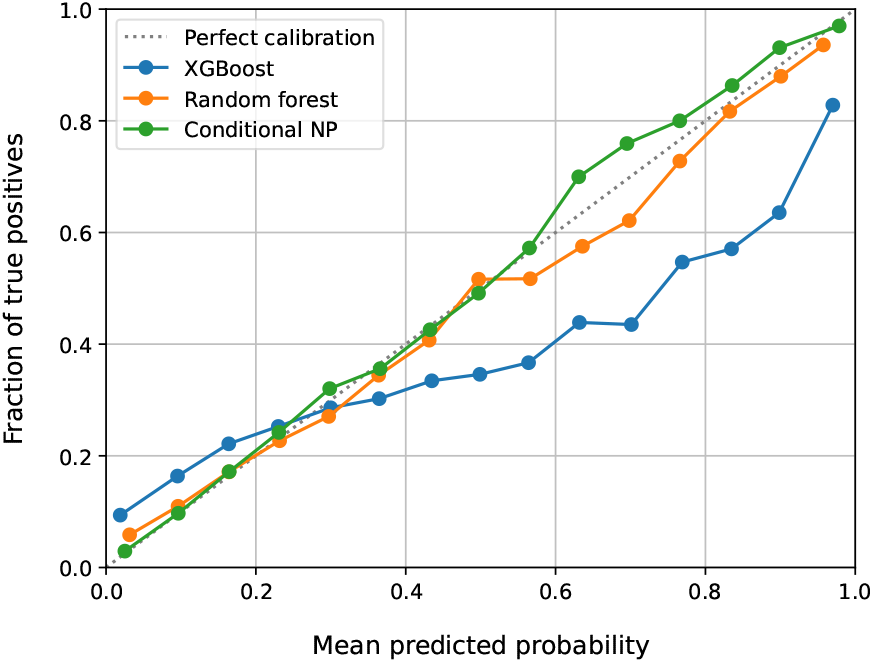
Calibration curve of CNP and baselines.

### 3.3 Performance across system organ classes

In our final experiment, we verified that the CNP produced accurate predictions for ADRs associated to any physiological system in SIDER. Achieving accurate ADR prediction across all organ classes is a desirable trait that would increase the usefulness of a model during medicinal development. Figure 3 shows the average balanced accuracy within different organ classes, evaluated on the functions and molecules in *ftest, dtest*. We included in our analysis every organ class represented by 3 or more *ftest* functions. As it can be seen, the CNP remains superior to baselines across all physiological systems.

**Figure 3:**
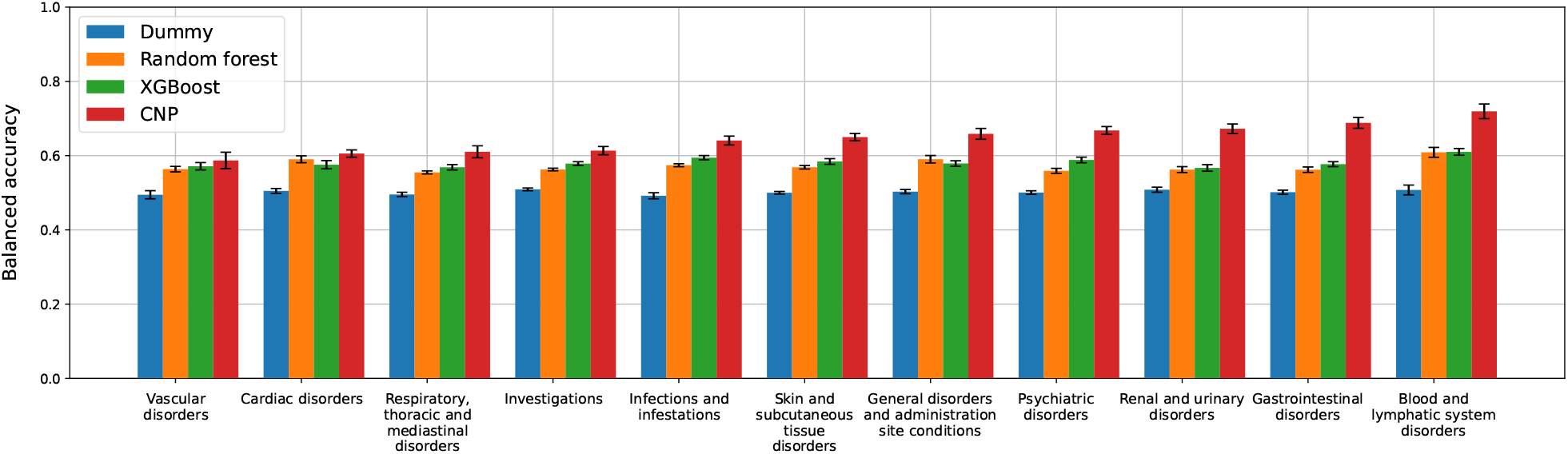
Balanced accuracy within physiological systems in SIDER. A classifier that always predicts the same class or one that predicts classes at random (blue bars) achieves a balanced accuracy 0.5.

## 4 Discussion

Our study suggests that a meta-learning approach to ADR prediction could bring significant benefits relative to single-task models typical in traditional chemoinformatic research. CNPs on molecular fingerprints showed more accurate and better calibrated classification of side effects than XGBoost or random forests on the same representations. We observed such improvements across a range of physiological systems.

It is worth noting that the exploratory evaluation presented in this work is limited to binary labels from SIDER, which neglects many complex features important for ADR such as severity, frequency, stratification of susceptible patient populations, dosage or drug interactions. These factors, which are clearly relevant for ADRs in the real world, are not always available in public datasets: for example, SIDER only has frequency information for 39.9% of its labels, and lacks details about any of the other aspects cited (Kuhn et al., 2016). Nonetheless, our study makes a compelling case for framing prediction of sparse ADR datasets as meta-learning. Further research is needed to explore the full potential of NPs in this area, as well as to perform more strict benchmarks against a transfer learning or multi-task approaches, in order to pave the way for their practical deployment in decision-making for medicinal chemists.

### Reproducibility statement

The SIDER dataset employed in this study can be accessed on http://sideeffects.embl.de/.

## References

Lucian Chan, Marcel Verdonk, and Carl Poelking. Embracing assay heterogeneity with neural processes for markedly improved bioactivity predictions, 2023.

Sanjoy Dey, Heng Luo, Achille Fokoue, Jianying Hu, and Ping Zhang. Predicting adverse drug reactions through interpretable deep learning framework. BMC Bioinf., 19(21):1–13, December 2018. ISSN 1471-2105. doi: 10.1186/s12859-018-2544-0.

Miguel Garcia-Ortegon, Andreas Bender, and Sergio Bacallado. Conditional neural processes for molecules, 2022.

Marta Garnelo, Dan Rosenbaum, Chris J. Maddison, Tiago Ramalho, David Saxton, Murray Shanahan, Yee Whye Teh, Danilo J. Rezende, and S. M. Ali Eslami. Conditional Neural Processes. arXiv, July 2018a. doi: 10.48550/arXiv.1807.01613.

Marta Garnelo, Jonathan Schwarz, Dan Rosenbaum, Fabio Viola, Danilo J. Rezende, S. M. Ali Eslami, and Yee Whye Teh. Neural Processes. arXiv, July 2018b. doi: 10.48550/arXiv.1807.01622.

Sunghwan Kim, Jie Chen, Tiejun Cheng, Asta Gindulyte, Jia He, Siqian He, Qingliang Li, Benjamin A. Shoemaker, Paul A. Thiessen, Bo Yu, Leonid Zaslavsky, Jian Zhang, and Evan E. Bolton. PubChem 2023 update. Nucleic Acids Res., 51(D1):D1373–D1380, January 2023. ISSN 0305-1048. doi: 10.1093/nar/gkac956.

Michael Kuhn, Ivica Letunic, Lars Juhl Jensen, and Peer Bork. The SIDER database of drugs and side effects. Nucleic Acids Res., 44(D1):1075–1079, January 2016. ISSN 1362-4962. doi: 10.1093/nar/gkv1075.

Chun Lee and YiPing Chen. Machine learning on adverse drug reactions for pharmacovigilance. Drug Discovery Today, 2019.

Eunjoo Lee, Jiho Yoo, Huisun Lee, and Seunghoon Hong. MetaDTA: Meta-learning-based drug-target binding affinity prediction. In ICLR2022 Machine Learning for Drug Discovery, 2022. URL https://openreview.net/forum?id=yzlif16IASM.

Ruoqi Liu and Ping Zhang. Towards early detection of adverse drug reactions: combining pre-clinical drug structures and post-market safety reports. BMC Med. Inf. Decis. Making, 19(1):1–9, December 2019. ISSN 1472-6947. doi: 10.1186/s12911-019-0999-1.

H. L. Morgan. The Generation of a Unique Machine Description for Chemical Structures-A Technique Developed at Chemical Abstracts Service. J. Chem. Doc., 5(2):107–113, May 1965. ISSN 0021-9576. doi: 10.1021/c160017a018.

Alexandru Niculescu-Mizil and Rich Caruana. Predicting good probabilities with supervised learning. In ICML ‘05: Proceedings of the 22nd international conference on Machine learning, pages 625–632. Association for Computing Machinery, New York, NY, USA, August 2005. ISBN 978-159593180. doi: 10.1145/1102351.1102430.

Igho J. Onakpoya, Carl J. Heneghan, and Jeffrey K. Aronson. Post-marketing withdrawal of 462 medicinal products because of adverse drug reactions: a systematic review of the world literature. BMC Med., 14(1):1–11, December 2016. ISSN 1741-7015. doi: 10.1186/s12916-016-0553-2.

F. Pedregosa, G. Varoquaux, A. Gramfort, V. Michel, B. Thirion, O. Grisel, M. Blondel, P. Prettenhofer, R. Weiss, V. Dubourg, J. Vanderplas, A. Passos, D. Cournapeau, M. Brucher, M. Perrot, and E. Duchesnay. Scikit-learn: Machine learning in Python. Journal of Machine Learning Research, 12:2825–2830, 2011.

RDKit. RDKit: Open-source cheminformatics. https://www.rdkit.org.

SIDER. SIDER dataset. Accessed on 29th September 2023. http://sideeffects.embl.de/.

Morgan Thomas, Andrew Boardman, Miguel Garcia-Ortegon, Yang Hongbin, Chris de Graaf, and Andreas Bender. Applications of Artificial Intelligence in Drug Design: Opportunities and Challenges. Methods Mol. Biol., 2390(1-59.):;, 2022. ISSN 1940-6029. doi: 10.1007/978-1-0716-1787-8_1.

David Weininger. SMILES, a chemical language and information system. 1. Introduction to methodology and encoding rules. J. Chem. Inf. Comput. Sci., 28(1):31–36, February 1988. ISSN 0095-2338. doi: 10.1021/ci00057a005.

XGB. XGBoost API. Accessed on 29th September 2023. https://xgboost.readthedocs.io/.

Fei Zhang, Bo Sun, Xiaolin Diao, Wei Zhao, and Ting Shu. Prediction of adverse drug reactions based on knowledge graph embedding. BMC Med. Inf. Decis. Making, 21(1): 1–11, December 2021. ISSN 1472-6947. doi: 10.1186/s12911-021-01402-3.

Wen Zhang, Xiang Yue, Feng Liu, Yanlin Chen, Shikui Tu, and Xining Zhang. A unified frame of predicting side effects of drugs by using linear neighborhood similarity. BMC Syst. Biol., 11(6):23–34, December 2017. ISSN 1752-0509. doi: 10.1186/s12918-017-0477-2.

